# Measuring the exposure of primate reservoir hosts to mosquito vectors in Malaysian Borneo

**DOI:** 10.1101/2021.05.25.445315

**Authors:** Rebecca Brown, Milena Salgado-Lynn, Amaziasizamoria Jumail, Cyrlen Jalius, Tock-Hing Chua, Indra Vythilingam, Heather M. Ferguson

## Abstract

Several vector-borne pathogens of primates have potential for human spillover. An example is the simian malaria *Plasmodium knowlesi* which is now a major public health problem in Malaysia. Characterization of exposure to mosquito vectors is essential for assessment of the force of infection within wild primate populations, however few methods exist to do so. Here we demonstrate the use of thermal imaging and Mosquito Magnet Independence Traps (MMIT) to assess the abundance, diversity and infection rates in mosquitoes host seeking near long-tailed macaque (*Macaca fasicularis*) sleeping sites in the Lower Kinabatangan Wildlife Sanctuary, Malaysian Borneo. The primary *Plasmodium knowlesi* vector, *Anopheles balabacensis*, was trapped at higher abundance near sleeping sites than control trees. Although none of the *An. balabacensis* collected (n=15) were positive for *P. knowlesi*, two were infected with another primate malaria *Plasmodium inui*. Analysis of macaque stools from sleeping sites confirmed a high prevalence of *Plasmodium* infection, suspected to be *P. inui. Plasmodium inui* infections have not yet been reported in humans, but its presence in *An. balabacensis* here and previously in human-biting collections highlight its potential for spillover. We advocate the use of MMITs for non-invasive sampling of mosquito vectors that host seek on wild primate populations.

## Introduction

Non-human primates (NHPs) are reservoirs of vector-borne pathogens that can infect humans. Some already pose significant public health problems, such as the monkey malaria parasites *Plasmodium simium* (1) and *P. brasilianum* (2) in South America. Several human vector borne diseases have sylvatic origins, including Yellow Fever, Zika and Dengue (3,4) alongside other lesser known viruses with potential to emerge (5). Understanding the force of infection in wild primate populations is crucial for assessment of potential for human spillovers, and the possibility of disrupting transmission in primate populations to mitigate against this. Such assessment requires estimating primate exposure to mosquito vectors, however there are few practical methods to do this. Current methods are invasive by relying on captive monkeys used as baits in traps, and may not reflect exposure in a natural population. There would be great value in finding a non-invasive and representative method for characterising NHP exposure to mosquito vectors.

One of the most notable primate Vector-borne diseases (VBD) of public health significance in SE Asia is *Plasmodium knowlesi*, a zoonotic malaria whose natural hosts are long-tailed and pig-tailed macaques and *Presbitis* leaf-monkeys, transmitted by Leucosphyrus group *Anopheles* (6–9). Since first detection of human cases in 2004 (10), *P. knowlesi* has become the most common cause of malaria in people in Malaysian Borneo (11). In 2014, human cases of another macaque malaria, *P. cynomolgi*, were also reported in Malaysia (12,13). Other macaque malarias (*P. coatneyi, P. fieldi* and *P. inui* (14,15)) have been detected in mosquito vectors in Malaysian Borneo, but are not yet known to have infected humans. Additional VBDs circulate in NHPs in Malaysia that can infect people; e.g. sylvatic Dengue in macaques, leaf monkeys and orangutans (5,16,17), and the filarial worm *Brugia malayi* in leaf monkeys. (18,19). This wide range of potential VBDs necessitates surveillance of mosquitoes biting NHP to anticipate future spillover risk.

Characterization of VBD transmission in primates has been hindered by logistical and ethical constraints. To date there are a limited range of tools for studying NHP exposure to VBDs; most being invasive by requiring blood sampling (20–22). Alternative non-invasive methods for detecting malaria parasite DNA in faecal samples (23–29) are promising but are yet to be widely applied and optimized. Similar constraints apply to assessment of primate exposure to mosquito vectors. This has generally been conducted through “Baited Traps” in which monkeys are placed in cages inside a net with gaps to allow mosquitoes attracted to enter but not leave (7,30–32). Contemporary animal welfare regulations for working with captive primates often make such approaches unfeasible. Alternative less invasive approaches such as “e-nets” in which macaques are held in larger cages and have their odour collected and channelled to attract mosquitoes are logistically challenging and yield few vectors (33). Finally, all methods that require the use of a host ‘bait’ require capture of wild primates or handling of captive individuals; both of which are invasive and could cause distress. Identification of less invasive methods for sampling the vector population that host seeks on wild primate populations would be of great value.

So far most investigation of the mosquito vectors of macaque VBDs have been conducted in areas near human settlements (30–33) which may not be reflective of natural transmission cycles within primate populations in the absence of humans. Knowledge of the range of parasite and vector species responsible for monkey to monkey transmission in undisturbed settings is necessary to identify future spillover risks, and assess the feasibility of disrupting transmission in primate reservoir populations as a strategy to reduce this.

Here we evaluated the use of commercially available Mosquito Magnet Independence Traps (MMIT) to passively sample malaria vectors host seeking in the vicinity of long-tailed macaques within the Lower Kinabatangan Wildlife Sanctuary (LKWS) Sabah, Malaysia. Aims were to assess the performance of the MMIT in terms of the abundance and diversity of potential vector species captured near macaque roosts versus uninhabited trees, and whether infection rates in vectors caught near roost sites were reflective of infection prevalence in macaques as assessed from faecal samples. We also investigated relationships between malaria vector abundance in MMITs, macaque troop size as measured with thermal imagery, and environmental factors. Whilst Mosquito Magnet traps have been investigated for passive surveillance of human malaria vectors (34–40), to our knowledge this is the first time it has been evaluated in a wild monkey population.

## Methods

### Study site

This study was conducted at the Danau Girang Field Centre (DGFC), Lot 6 of the LKWS (5°24′49.93″ N, 118°02′18.58″ E) (Fig. 1). The LKWS is a protected secondary disturbed forest area (ranging from 10 - 60 years old) that contains primary to secondary lowland dipterocarp forest, mangrove and oil palm plantations (41,42). The sanctuary spans 27000ha (43), and hosts ten primate species including the two reservoir hosts of *P. knowlesi*; long-tailed macaques (*Macaca fasicularis*), and pig-tailed macaques (*Macaca nemestrina*). In 2002, population densities (per km2) were estimated as 16.82 for *M. fasicularis* and 3.30 for *M. nemestrina* (36). The nearest human settlement is at least 15 km downstream from DGFC.

**Figure 1.**
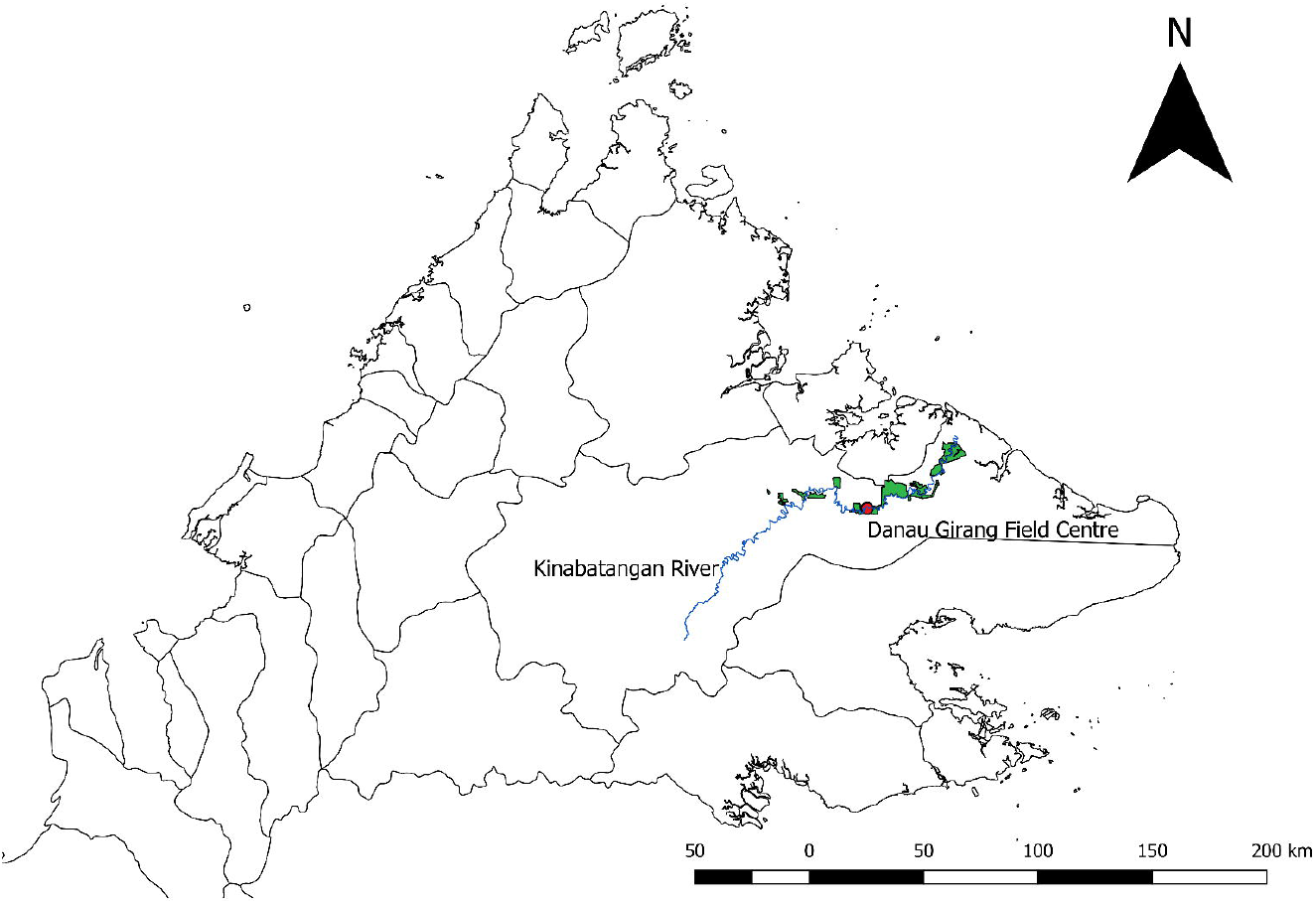
Map of Sabah indicating the location of the Danau Girang Field Centre (red) along the Kinabatangan river (blue). Green areas indicate boundaries of the Lower Kinabatangan Wildlife Sanctuary (Lots 1 - 10) and black lines show administrative districts.

### HLC vs MMIT trap comparison

A pilot study was performed to confirm the MMIT (Mosquito Magnet, model: MM3200, supplier: Syarikat Thiam Siong Sdn Bhd, Sabah) was capable of sampling *Anopheles* in this environment (Fig. S1). The MMIT was compared with the standard Human Landing Catch (HLC) method; which is known to be efficient for sampling the *P. knowlesi* vector *An. balabacensis* (14,30,32). The MMIT lures mosquitoes using mammalian odour bait (CO_2_ and octenol), heat and water vapour (36,39). The MMIT was modified to run off batteries (4 x 1.5V) and on locally available gas (30% propane: 70% butane).

Each night, one HLC and one MMIT site were selected; with stations ~100m apart on one of three walking trails (Fig. S2). The following night the HLC and MMIT switched sites in a cross over design. This was repeated for ten nights of collections. Hourly collections were conducted from 18:00 – 00:00 hrs to coincide with the peak biting time of *An. balabacensis* (18:00 - 20:00 hrs (14,44)). One person performed HLC accompanied by an assistant. Each hour comprised 45 minutes of trapping and 15 minutes break. The MMIT was switched off during the break and the collection net replaced.

### Use of MMIT for sampling vectors near macaque sleeping sites

Mosquito sampling using MMIT was conducted along a 20km section of the Kinabatangan river (560km, Fig. 1). Trees (*Colona, Nauclea subdita, Pterospermum acerifolium, Kleinhovia hospita* and *Ficus*) of a minimum 20m depth lining the river bank (45) are used as sleeping sites for several primate species (46,47), including long-tailed macaques. The study site was divided into ten 2km transects (Fig. S3). The home range of long-tailed macaques in this reserve was estimated as 1.25km^2^ in a previous survey (47). To avoid repeated sampling near the same macaque troop, sampling was conducted in different 2km transects each night. Mosquito sampling took place in each transect once every ten nights; with the transect selected randomly using the Random UX app. Sampling was conducted in blocks of five nights with a one night break; resulting in 38 sampling nights between September to November of 2017. Each block was sampled 3 - 4 times during this period, with traps placed on alternate sides of the river on each visit.

Upon arrival at the selected transect (17:30 hrs), the river banks were scanned with a thermal imaging camera to identify potential macaque troops by driving slowly up and down the river. When the camera indicated presence of a troop, binoculars were used to inspect trees for long-tailed macaques. Once the presence of roosting macaques was confirmed, a MMIT was placed near the bottom of their sleeping tree (conditional on bank being accessible, Fig. S4). Macaques generally moved from the selected tree to higher up in the canopy or deeper inside the forest as the boat approached, but would return after the trap was placed. A second MMIT was placed at least 100m away at a ‘control’ tree that was of similar structure and species, but uninhabited by macaques or other primates. This ‘control’ tree enabled differentiation of mosquitoes specifically host seeking in the vicinity of macaques.

Mosquitoes were collected at macaque sleeping and control sites each night from 18:00 – 06:00 hrs. Before sunrise and movement of macaques (approximately 05:30 hrs), the number of macaques sleeping in the tree where the MMIT was placed were counted from the boat using the thermal camera. Daily rainfall data (collected by rain gauge) was provided by DGFC.

### Mosquito processing

Mosquitoes were stored at −20°C for approximately 12 hrs then identified to genera and species where possible (48–51). Leucosphyrus group *Anopheles* were identified using Sallum *et al* (2005) (52). All identified mosquitoes were stored in 95% ethanol. Molecular analysis was performed on Leucosphyrus group *Anopheles, An. barbirostris gp*. (*An. barbirostris and An. donaldi*), *An. epiroticus and An. tesselatus* (malaria vectors in Sabah or elsewhere in SE Asia, (44,50,53–56)) to screen for *Plasmodium* infections using the method described in (57).

### Macaque Faecal collection

Each morning after emptying MMIT traps, the ground within a 20m radius of sleeping trees was inspected for the presence of fresh macaque stools (see Supplementary Methods S1). Stool samples were homogenized in RNAlater solution then stored at −20°C.

DNA was extracted from 200 μl of each stool solution using the QIAamp DNA Stool Mini Kit. DNA was eluted in 100 μl buffer AE and stored at −20 °C. Samples were screened by PCR for detection of DNA from the *Plasmodium* genus (see Supplementary Methods S2). *Plasmodium* positive samples were then screened to test for the specific presence of *P. knowlesi* following the method of Kawai *et al* (29).

### Statistical analysis

Data were analysed using R statistical programming software (3.4.2) with packages lme4 and multcomp. Generalized Linear Mixed Models (GLMMs) were used to compare the abundance of mosquitoes in HLC and MMIT; with comparisons made for all mosquitoes and just *Anopheles*.Negative binomial GLMMs were used to account for overdispersion in mosquito count data (58). The response variable was the mean abundance of (i) all mosquitoes (ii) *Anopheles* per night. The main fixed effect was trap type with random effects fit for date and trail. A post hoc Tukeys’ test was used to assess differences in mosquito abundances between traps. The vegan package was used to measure *Anopheles* diversity in HLC and MMIT catches. Four diversity indices were calculated: species richness, rarefied species richness, Simpson’s index and the Shannon index (57).

Sampling of mosquitoes near trees where macaques were sleeping was conducted for 38 nights. On a few occasions, macaques or other primates were present at the control site in the mornings or the traps stopped working overnight due to failure of gas supply or batteries. Excluding these scenarios, data was available from 33 nights of sampling at control trees and 34 nights at trees with sleeping macaques. With this data, GLMMs were constructed to test for differences in the response variables of 1) *Anopheles* abundance 2) *An. balabacensis* abundance and 3) *An. donaldi* abundance between macaque sleeping sites and control trees. A negative binomial distribution was used with date and river transect set as random effects. Models tested for associations between mosquito abundance and macaque presence and abundance, and rainfall on the day of sampling. The significance of each variable was tested by backward elimination using likelihood ratio tests. Post-hoc Tukey’s tests were performed to assess differences in mosquito abundance between sleeping sites and control collections.

On nine sampling nights, the datalogger failed to record the average nightly temperature. Thus a subset of data (28 nights at control trees and 29 nights at sleeping sites) for which mean nightly temperatures were available was used to test for associations between temperature and mosquito abundance. Here the impact of temperature on three mosquito subgroups were investigated as above.

## Results

### HLC vs MMIT trap comparison

Both HLC and MMITs collected mosquitoes belonging to the same eight genera (Table S1). Mosquitoes were identified to species level where possible, however due to time constraints, priority was given to *Anopheles, Culex* and *Mansonia. Aedes* and *Uranotaenia* mosquitoes were mostly identified to subgenus. In general, mosquitoes trapped by HLC were in better condition for morphological identification than those trapped in the MMIT because key characteristics necessary for species determination such as hairs and scales were better preserved.

Almost all *Anopheles* caught in the HLC could be speciated, except one individual that was missing features to distinguish between *An. barbirostris* or *An. donaldi*. Two *Anopheles* from MMIT collections (3.2 % of total) could not be placed to a subgenus. Five *Anopheles* species were collected by HLC compared to 8 species with MMIT (Table 1). *Anopheles* diversity was higher in MMIT than HLC collections (Table 2). Both methods trapped the *P. knowlesi* vectors *An. balabacensis* and *An. donaldi*; with a higher proportion of these being caught by HLC (80.5 %, n = 29) than MMIT (72.6 %, n = 45).

**Table 1.**
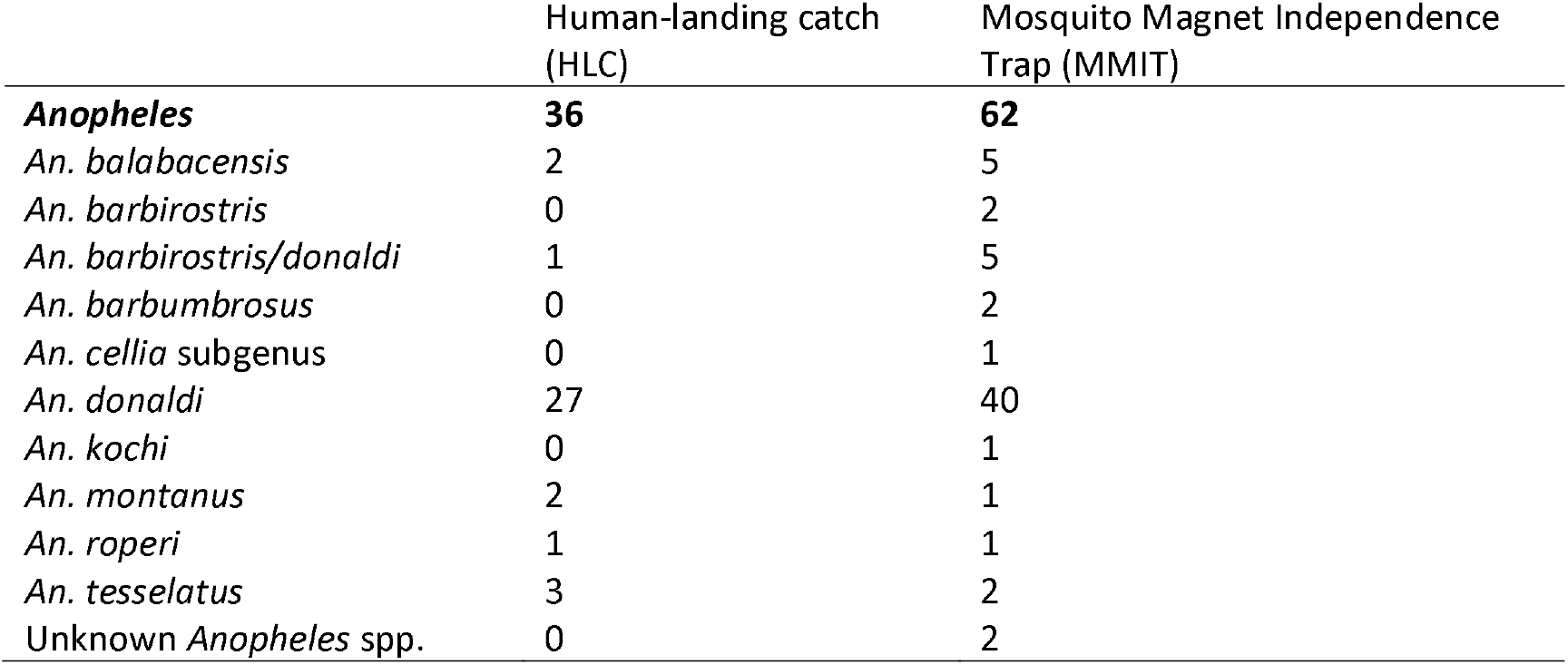
*Anopheles* mosquitoes caught by Mosquito Magnet Independence Trap (MMIT) and human-landing catch (HLC) over ten nights of trap comparison study in Lower Kinabatangan Wildlife Sanctuary, Sabah.

**Table 2.**
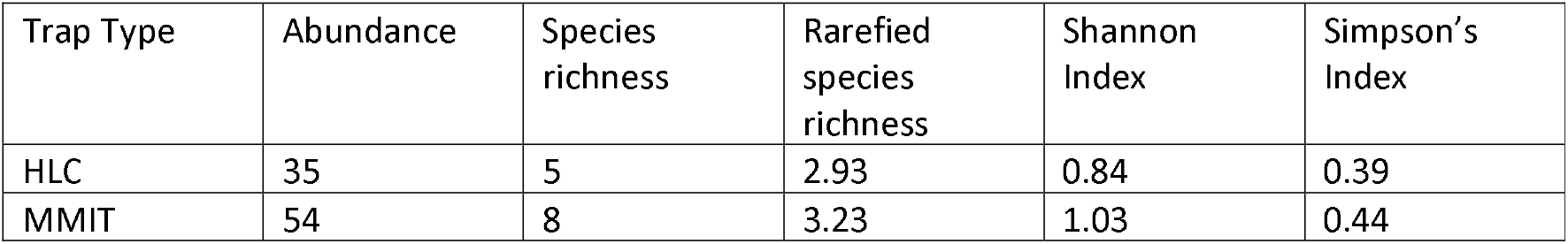
Measures of diversity in *Anopheles* species from Mosquito Magnet Independence Trap (MMIT) and human-landing catch (HLC) collections from a ten-night trap comparison study in Lower Kinabatangan Wildlife Sanctuary, Sabah.

| Trap Type | Abundance | Species richness | Rarefied species richness | Shannon Index | Simpson's Index |
| --- | --- | --- | --- | --- | --- |
| HLC | 35 | 5 | 2.93 | 0.84 | 0.39 |
| MMIT | 54 | 8 | 3.23 | 1.03 | 0.44 |

Although mosquito numbers tended higher in MMIT than HLC collections, the mean nightly abundance was not significantly different (Tukey’s test: *P* = 0.39, Fig. 2A). Similarly, the mean nightly abundance of *Anopheles* did not vary between trapping methods (Tukey’s test: *P* = 0.210, Fig. 2A). *Anopheles donaldi* was most abundant between 18:00 – 20:00 hrs (Fig. 2C), whereas *An. balabacensis* biting rates were relatively constant between 18:00 - 23:00 hrs with none collected between 23:00 - 24:00 hrs (Fig. 2B).

**Figure 2.**
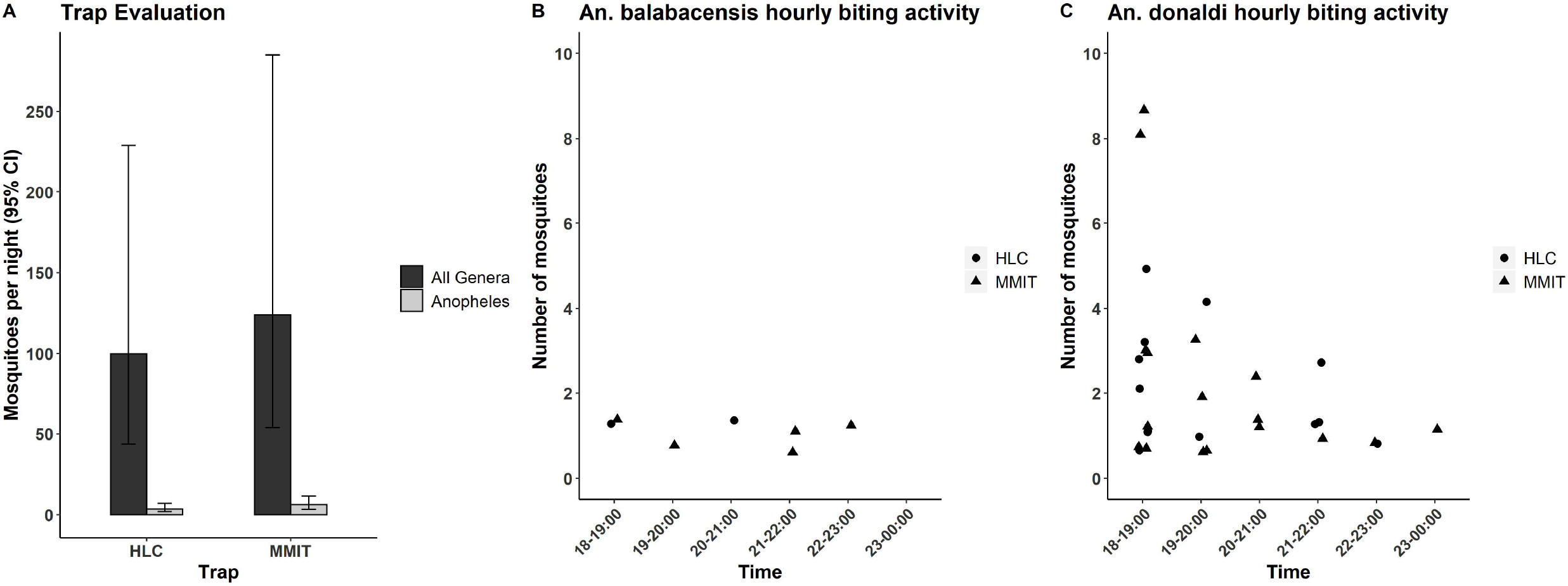
A) Mean abundance of mosquitoes caught per night by Human landing catch (HLC) and Mosquito Magnet Independence Trap (MMIT) as predicted by negative binomial generalised linear mixed models (GLMM). Error bars represent 95% confidence intervals B) *An. balabacensis* and C) *An. donaldi* trapped per hour by human-landing catch (HLC) and Mosquito Magnet Independence Trap (MMIT).

### MMIT to sample Anopheles host seeking near macaques

Overall, 11400 mosquitoes from eight genera were collected in MMITs placed near macaque sleeping sites and control trees (Table S2). Both malaria vector species, *An. balabacensis* and *An. donaldi* were trapped at sleeping sites and control trees.

Combining over all species in the genera, the mean nightly abundance of *Anopheles spp*. was not significantly associated with the presence (X^2^ = 0.45, *df* = 1, *P* = 0.50) or number of macaques (X^2^ = 0.62, *df* = 1, *P* = 0.43) at a tree, or with daily rainfall (X^2^ = 0.67, *df* = 1, *P* = 0.41) (Fig. S5). Within the subset of data for which full temperature recordings were available, a strong positive association was detected between mean nightly *Anopheles* abundance and temperature (X^2^ = 6.46, *df* = 1, *P* =0.01, Fig. 3).

**Figure 3.**
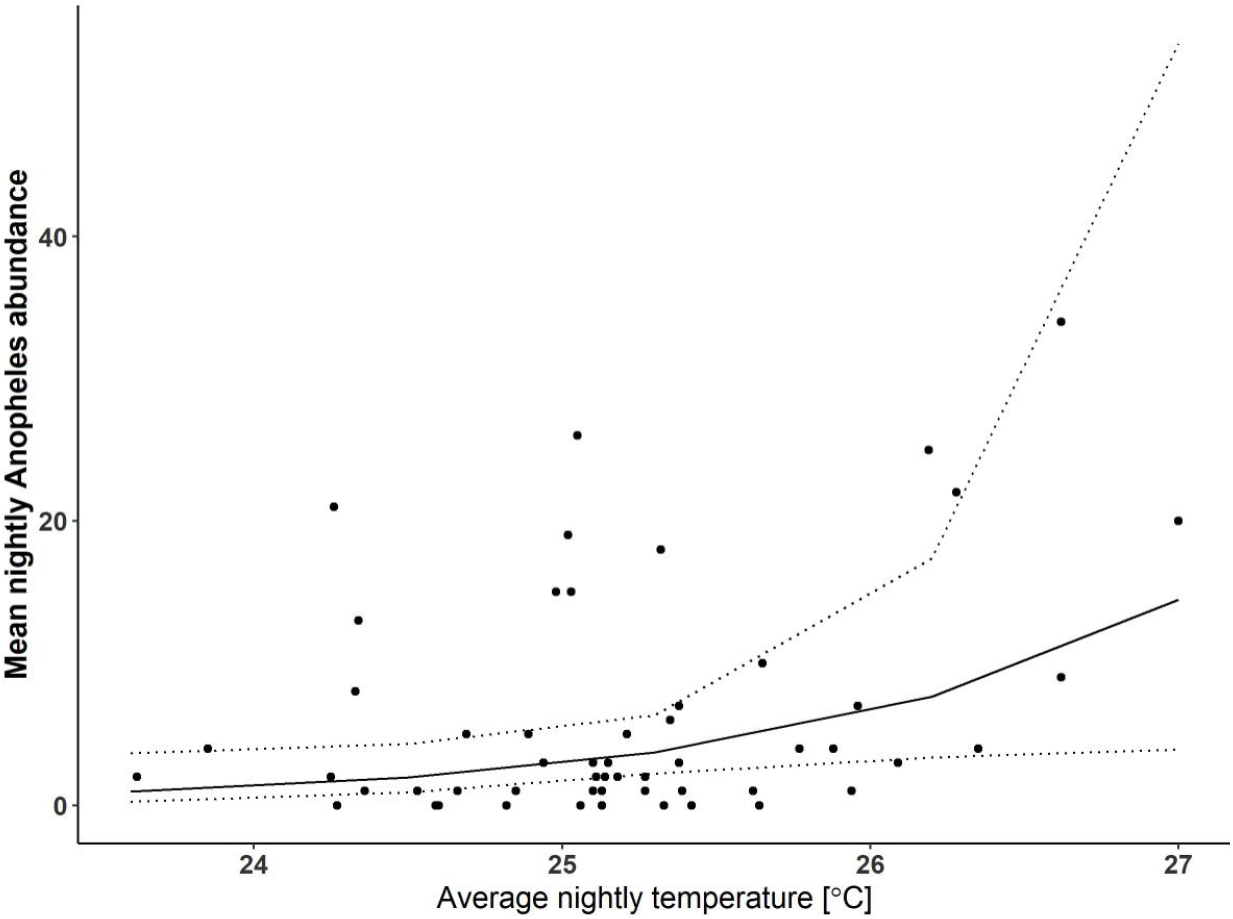
Predicted relationship between mean *Anopheles* abundance collected by Mosquito Magnet Independence Traps (MMIT) and average nightly temperature (from the subset of 28 sampling nights at control trees and 29 sampling nights at macaque sleeping sites for which environmental data were available). Points indicate observed data, with the line indicating the predicted association. Dashed lines represent upper and lower 95% confidence intervals.

The primate malaria vector *An. balabacensis*, however, was significantly impacted by the presence of macaques at sampling sites. The mean abundance of *An. balabacensis* was significantly higher near macaque roost sites than at control trees ((X^2^ = 8.25, *df* = 1, *P* < 0.01, Fig. 4A), but was not related to number present (X^2^ = 1.28, *df* = 1, *P* = 0.26) or daily rainfall (X^2^ = 0.42, *df* = 1, *P* = 0.52, Fig. 4B and C). Sample sizes of *An. balabacensis* within the subset of data where temperature was available were not sufficient for robust analysis.

**Figure 4.**
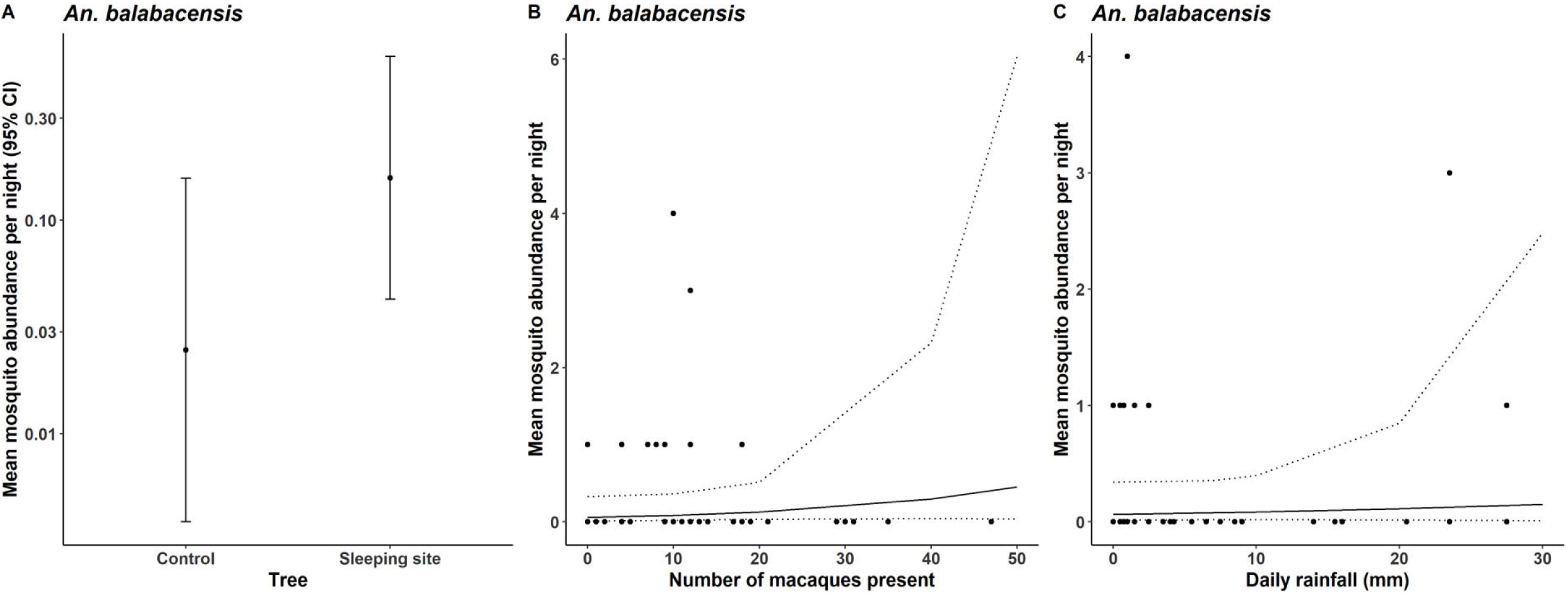
Influence of A) macaque presence/absence, B) number of macaques present and C) daily rainfall on the mean nightly *An. balabacensis* abundance collected by Mosquito Magnet Independence Traps (MMIT). Points indicate observed data in B and C, with the line indicating the predicted association. Error bars and dashed lines are 95% confidence intervals and * represents *P* < 0.05.

The total number of *An. donaldi*, caught at macaque sleeping sites (n = 106) was lower than at control trees (n = 211, Table 3); however this difference was only marginally significant (Tukeys: *P* = 0.052). The abundance of *An. donaldi* was not dependent on the presence or absence of macaques (X^2^ = 0.92, *df* = 1, *P* = 0.34), the number of macaques present (X^2^ = 0.13, *df* = 1, *P* = 0.72) or the daily rainfall (X^2^ = 0.23, *df* = 1, *P* = 0.63) (Figure S6). Within the subset of data for which full temperature recordings were available, *An. donaldi* abundance was positively associated with mean daily temperature (X^2^ = 5.86, *df* = 1, *P* = 0.02,Figure S7).

**Table 3.**
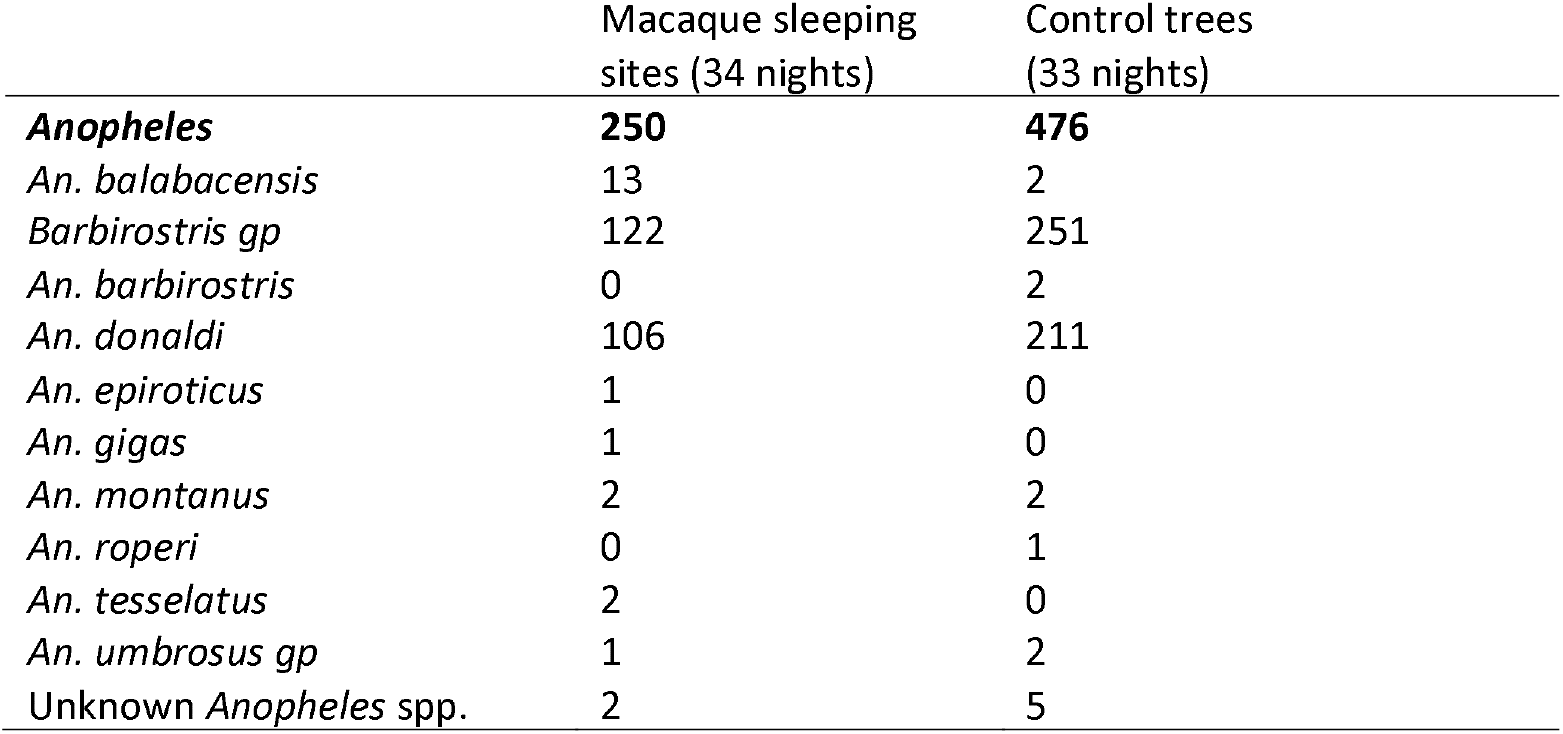
*Anopheles* mosquitoes caught with Mosquito Magnet Independence Trap (MMIT) at trees with and without sleeping macaques (control trees) within the Lower Kinabatangan Wildlife Sanctuary, Sabah.

### Plasmodium infections in mosquitoes and macaque stools

Eighty-one *Anopheles* collected in the initial HLC versus MMIT trap comparison were tested for malaria (*An. donaldi* = 61, *An. balabacensis* = 7, *An. barbirostris/donaldi* = 5, *An. tesselatus* = 5, *An. celia* group = 1, *An*. unknown = 2). Of these, one tested positive for *Plasmodium* infection (n = 1/81). In the larger study using MMITs at macaque sleeping sites and control trees, 398 *Anopheles* were tested for malaria (Barbirostris group = 373 (including *An. barbirostris* (2) and *An. donaldi* (317)), *An. balabacensis* = 15, *An. epiroticus* = 1, *An. tesselatus* = 2 and unidenitifed *Anopheles* species = 7, Table 3). Of these, one tested positive for *Plasmodium* (n = 1/398). Both infections were confirmed to be *P. inui*. Both infections were detected in *An. balabacensis*, representing an overall infection rate of 9% (n = 2/22) in this vector species.

Of the 46 long-tailed macaque faecal samples collected, 17 (37%) tested positive for *Plasmodium*.However in the subsequent round of PCR analysis to test for *P. knowlesi*, none were positive. Samples were not screened for other malaria species, thus the identity of *Plasmodium* infections remains unknown.

## Discussion and Conclusions

This study to our knowledge provides the first evaluation of the use of MMIT for sampling the mosquitoes that host seek on wild primates. We show that the abundance of all mosquitoes (pooled across genera), and *Anopheles* in particular was similar in collections made by MMITs and the HLC gold standard. Our results confirm that the MMIT can be used as an indirect exposure-free alternative to the HLC. MMIT placed below macaque roosting trees collected several known vectors of zoonotic and human malaria (*An. balabacensis, An. donaldi* and *An. barbirostris*). While *Anopheles* density was not higher overall at trees with than without macaques; the abundance of the confirmed primate vector *An. balabacensis* was significantly higher near macaque sleeping sites. This implies *An. balabacensis* were actively host seeking on macaques. Macaque density was very high in the study area, with analysis of their faecal samples indicating a high prevalence of *Plasmodium* infection (37%). However, the zoonotic malaria *P. knowlesi* was not detected in either vector or macaque samples here; suggesting transmission was primarily of other simian parasite species. Mosquito infections were confirmed to be *P. inui*; a primate malaria parasite that has not been documented in humans. Despite the absence of *P. knowlesi*, the ability of MMITs to detect known vector species feeding at macaque sleeping sites highlights its value for non-invasive monitoring of primate exposure to mosquito vectors. This tool could thus provide opportunity to study the transmission of *P. knowlesi* as well as other primate VBDs in wild primate populations.

The abundance of *Anopheles spp*. was similar in MMIT and HLC collections, although the MMIT caught more species (8 vs 5 in the HLC). The greater diversity of Anophelines in the MMIT compared to HLCs, also seen in a Venezuelan setting (59) may be due to the use of a general R-octenol bait that attracts both anthropophilc and zoophilic mosquitoes (60), and/or that it releases a higher concentration of host cues than a single human collector. Here only one individual performed HLC, but it is well known that volatile emissions vary between people (61). Further investigation using multiple participants in HLC is required for more robust evaluation of the relative performance of MMIT and HLC methods. However, given the MMIT has an advantage of enabling passive sampling and collected just as many *Anopheles* as HLC, it may be a more practical approach for sampling of malaria vectors host seeking on wild primates.

The primary *P. knowlesi* vector in Sabah, *An. balabacensis*, was detected at significantly higher numbers near trees with than without macaques; indicating this species has a propensity for this host type. This finding is in line with a host choice study involving baited electrocuting nets where *An. balabacensis* fed on both monkey and human hosts (33). However, there was no significant relationship between *An. balabacensis* abundance and the number of macaques at the sleeping site. Macaque troop size varied across sampling nights from 2 – 47 individuals (average ~14), thus incorporating substantial variability for detecting an association with mosquito density. Studies examining the influence of host density on malaria vector density have found correlation (positive and negative depending on vector species) between adult Anopheline density and the density of humans (62,63). The lack of association here, however may be the result of the odour plume of even one macaque being sufficient to lure *An. balabacensis*. Alternatively, mosquitoes could be attracted to the trees themselves. Macaques are known to revisit sleeping trees (64) thus macaque odour cues could build up around a site, signalling a reliable bloodmeal source for vectors. Additionally, there could be environmental characteristics not measured here that contributed to higher *An. balabacensis* abundances at macaque sleeping sites.

Associations between *Anopheles* abundance in MMIT collections and rainfall and temperature were not detected. Higher temperature and rainfall has been demonstrated to increase mosquito abundances in MMITs used in the Brazilian rainforest (38). No association was detected between temperature or rainfall and *An. balabacensis*; but ability to test for this was limited by small sample sizes. To investigate seasonal fluctuations in vector abundance with rainfall and temperature, we recommend more intensive longitudinal sampling across a full year to increase sample sizes and capture the extremes of environmental variation.

The primary focus of this study was investigation of *P. knowlesi* transmission within its wildlife reservoir in the absence of humans. However, malaria infections were detected in only two *An. balabacensis* and in both cases it was *P. inui. Plasmodium inui* is commonly found in wild macaques (65) and can infect humans under experimental conditions (via blood transfusion or infected mosquito bites in the lab (66). However natural human infections of *P. inui* have not been documented. Relatively high rates of *P. inui* and other primate malarias (*P. cynomolgi, P. fieldi and P. coatneyi*) have been described in *An. balabacensis* in village settings in Sabah (15). Thus, people are regularly exposed to these parasites in peri-domestic as well as forest settings; raising the possibility that *P. inui* could pose a significant risk for zoonotic spillover in the future.

The absence of *P. knowlesi* infection in mosquito vectors was matched with its absence in the macaque population. Although *Plasmodium* DNA was detected in more than a third of macaque stool samples, but none of these samples were identified as *P. knowlesi*. We hypothesise these infections were most likely *P. inui* based on its confirmation in the local *An. balabacensis* population. *Plasmodium knowlesi* prevalence in macaques has been reported at 6.9 % and 30 % in Peninsular Malaysia (31,67), and 20 % and 86.6 % in Sarawak (68,69). However these estimates were derived from analysis of macaque blood samples which have greater sensitivity to detect low density infections than the faecal screening method used here (70). However, another study also based on analysis of macaque blood samples reported a much lower prevalence of *P. knowlesi* (0.4 % (71)); indicating macaque infection rates are naturally heterogeneous. Researchers often assume that the force of *P. knowlesi* infection coming from macaques to humans is high throughout Sabah; given one study found P. knowlesi infection in 20% of the faeces collected from wild long-tailed macaques within the Kudat District, the hotspot of human infection in 2013 - 2014 (Salgado-Lynn, unpublished data). In the same study, 80% of the blood samples of macaques from a different district in Sabah were positive for *Plasmodium*, 66% of which were positive for *P. knowlesi*. However the apparent absence of *P. knowlesi* infection here indicates the force of infection may vary considerably between wild reservoir populations. Therefore, recent efforts to generate *P. knowlesi* risk maps based on macaque distribution (79–81) may be limited by failure to incorporate underlying variation in infection prevalence within macaque populations. Furthermore, blanket control policies based on macaque culling may be both ethically questionable and have limited impact.

Here we demonstrate the suitability of MMIT for sampling mosquitoes host seeking in the vicinity of macaques and advocate its use as a tool for monitoring vector borne infections circulating in wild primate populations. In addition to its use for investigation into vector ecology associated with macaque to macaque pathogen transmission, it is a reliable alternative to performing HLC to study vectors feeding on people and removes the need to expose volunteers to potentially infectious mosquito bites. As yet there are no records of natural *P. inui* infections in people in Malaysia, however with the detection of the parasite in *An. balabacensis* here and in this vector species from collections nearby homes (15,72,73) people are likely frequently being exposed to this parasite and therefore warrants close surveillance of its potential for spillover.

## List of abbreviations

HLC: Human Landing Catch
MMIT: Mosquito Magnet Independence Trap
LKWS: Lower Kinabatangan Wildlife Sanctuary
GLMM: Generalised Linear Mixed Models

## List of Supplementary Files

Figure S1. A Mosquito Magnet Independence Trap (MMIT, left) and a view of the mosquito collection net within the MMIT (right).

Figure S2. Map of forest trails surrounding the main building of Danau Girang Field Centre in the Lower Kinabatangan Wildlife Sanctuary. Boxes indicate the sites used on Ficus (wet lowland forest), Kingfisher (wet lowland forest) and Kayu Malam (dry lowland forest) walking trails for human-landing catch (HLC) and Mosquito Magnet Independence Trap (MMIT) evaluation of collecting *Anopheles*. Blue lines depict bodies of water.

Figure S3. The 20km stretch of the Kinabatangan River surrounding Danau Girang Field Centre where macaque roosting sites and control trees were selected for mosquito collection. Purple dots indicate the boundary of each 2km transect that could be randomly selected for mosquito sampling on each night.

Figure S4. The Mosquito Magnet Independence Trap (MMIT) in position on the riverbank at the base of a *Ficus* (fig) tree to be used by a long-tailed macaque troop as their overnight resting place.

Figure S5. Influence of A) macaque presence/absence, B) number of macaques present and C) daily rainfall on the mean nightly *Anopheles* abundance collected by Mosquito Magnet Independence Traps (MMIT). Points indicate observed data in B and C, with the line indicating the predicted association. Error bars and dashed lines are 95% confidence intervals and * represents *P* < 0.05.

Figure S6. Influence of A) macaque presence/absence, B) number of macaques present and C) daily rainfall on the mean nightly *An. donaldi* abundance collected by Mosquito Magnet Independence Traps (MMIT). Points indicate observed data in B and C, with the line indicating the predicted association. Error bars and dashed lines are 95% confidence intervals and * represents *P* < 0.05.

Figure S7. Predicted relationship between mean *An. donaldi* abundance collected by Mosquito Magnet Independence Traps (MMIT) and average nightly temperature (from the subset of 28 sampling nights at control trees and 29 sampling nights at macaque sleeping sites for which environmental data were available). Points indicate observed data, with the line indicating the predicted association. Dashed lines represent upper and lower 95% confidence intervals.

Table S1. Mosquitoes caught by Mosquito Magnet Independence Trap (MMIT) and human-landing catch (HLC) over ten nights of trap comparison study in Lower Kinabatangan Wildlife Sanctuary, Sabah.

Table S2. Mosquitoes caught with Mosquito Magnet Independence Trap (MMIT) at trees with and without sleeping macaques (control trees) within the Lower Kinabatangan Wildlife Sanctuary, Sabah.

## Notes

https://doi.org/10.7910/DVN/ZS4VRY

### Competing Interest Statement

The authors have declared no competing interest.

## References

1. Brasil P, Zalis MG, Pina-costa A De, Siqueira AM, Júnior CB, Silva S. Outbreak of human malaria caused by Plasmodium simium in the Atlantic Forest in Rio de Janeiro: a molecular epidemiological investigation. Lancet. 2017;

2. Lalremruata A, Magris M, Vivas-Martinez S, Koehler M, Esen M, Kempaiah P, et al. Natural infection of Plasmodium brasilianum in humans: Man and monkey share quartan malaria parasites in the Venezuelan Amazon. EBioMedicine. 2015;2(9): 1186–92.

3. Rodhain F. The role of monkeys in the biology of dengue and yellow fever. Comp Immunol Microbiol Infect Dis. 1991;14(1):9–19.

4. Vorou R. Zika virus, vectors, reservoirs, amplifying hosts, and their potential to spread worldwide: What we know and what we should investigate urgently. Int J Infect Dis [Internet]. 2016;48:85–90. Available from: http://dx.doi.org/10.1016/j.ijid.2016.05.014

5. Valentine MJ, Murdock CC, Kelly PJ. Sylvatic cycles of arboviruses in non ⍰human primates. Parasit Vectors. 2019;12(463):1–18.

6. Knowles RM. A study of monkey malaria and its experimental transmission to man. Indian Med Gaz. 1932;67:301–20.

7. Wharton RH, Eyles D. E, Warren M. The development of methods for trapping the vectors of monkey malaria. Ann Trop Med Parasitol. 1963;57(I):32–46.

8. Warren M, Wharton RH. The vectors of simian malaria: identity, biology, and geographical distribution. J Parasitol. 1963;49(6):892–904.

9. Jeyaprakasam NK, Liew JWK, Low VL, Wan-Sulaiman WY, Vythilingam I. Plasmodium knowlesi infecting humans in Southeast Asia: What’s next? PLoS Negl Trop Dis [Internet]. 2020;14(12):e0008900. Available from: http://dx.doi.org/10.1371/journal.pntd.0008900

10. Singh B, Kim Sung L, Matusop A, Radhakrishnan A, Shamsul SSG, Cox-Singh J, et al. A large focus of naturally acquired Plasmodium knowlesi infections in human beings. Lancet. 2004 Mar;363(9414): 1017–24.

11. Hussin N, Lim YAL, Goh PP, William T, Jelip J, Mudin RN. Updates on malaria incidence and profile in Malaysia from 2013 to 2017. Malar J [Internet]. 2020; 19(1): 1–14. Available from: https://doi.org/10.1186/s12936-020-3135-x

12. Ta TH, Hisam S, Lanza M, Jiram AI, Ismail N, Rubio JM. First case of a naturally acquired human infection with Plasmodium cynomolgi. Malar J. 2014;13(1):1–7.

13. Law Y-H. Rare human outbreak of monkey malaria detected in Malaysia. Nature news. 2018.

14. Wong ML, Chua TH, Leong CS, Khaw LT, Fornace K. Seasonal and spatial dynamics of the primary vector of Plasmodium knowlesi within a major transmission focus in Sabah, Malaysia. PLoS Negl Trop Dis. 2015;9(10):1–15.

15. Manin BO, Ferguson HM, Vythilingam I, Fornace K, William T, Torr SJ, et al. Investigating the Contribution of Peri-domestic Transmission to Risk of Zoonotic Malaria Infection in Humans. PLoS Negl Trop Dis. 2016;10(10):1–14.

16. Rossi SL, Nasar F, Cardosa J, Mayer S V., Tesh RB, Hanley KA, et al. Genetic and phenotypic characterization of sylvatic dengue virus type 4 strains. Virology. 2012;423(1):58–67.

17. Young KI, Mundis S, Widen SG, Wood TG, Tesh RB, Cardosa J, et al. Abundance and distribution of sylvatic dengue virus vectors in three different land cover types in Sarawak, Malaysian Borneo. Parasites and Vectors. 2017;10(1):1–14.

18. Kwa BH. Environmental change, development and vectorborne disease: Malaysia’s experience with filariasis, scrub typhus and dengue. Environ Dev Sustain. 2008;10(2):209–17.

19. Cheong WH, Loong KP, Mahadevan S, Mak JW, Kan SK. Mosquito fauna of the Bengkoka Peninsula, Sabah, Malaysia. Southeast Asian J Trop Med Public Health. 1984; 15(1): 19–26.

20. Martinelli A, Culleton R. Non-human primate malaria parasites: Out of the forest and into the laboratory. Parasitology. 2018;145(1):41–54.

21. Deane L. Monkey malaria in Brazil-a summary of studies performed in 1964-1966. Geneva; 1967.

22. Dissanaike AS. Simian Malaria Parasites of Ceylon. Bull World Health Organ. 1965;32(1910):593–7.

23. Liu WM, Li YY, Learn GH, Rudicell RS, Robertson JD, Keele BF, et al. Origin of the human malaria parasite Plasmodium falciparum in gorillas. Nature. 2010;467(7314):420–U67.

24. Mapua MI, Petrželková KJ, Burgunder J, Dadáková E, Brožová K, Hrazdilová K, et al. A comparative molecular survey of malaria prevalence among Eastern chimpanzee populations in Issa Valley (Tanzania) and Kalinzu (Uganda). Malar J. 2016;15(1):1–11.

25. De Nys HM, Calvignac-Spencer S, Thiesen U, Boesch C, Wittig RM, Mundry R, et al. Age-related effects on malaria parasite infection in wild chimpanzees. Biol Lett. 2013;9(4).

26. de Assis GMP, de Alvarenga DAM, Costa DC, de Souza Junior JC, Hirano ZMB, Kano FS, et al. Detection of Plasmodium in faeces of the new world primate Alouatta clamitans. Mem Inst Oswaldo Cruz. 2016;111(9):570–6.

27. Abkallo HM, Liu W, Hokama S, Ferreira PE, Nakazawa S, Maeno Y, et al. DNA from pre-erythrocytic stage malaria parasites is detectable by PCR in the faeces and blood of hosts. Int J Parasitol [Internet]. 2014;44(7):467–73. Available from: http://dx.doi.org/10.1016/j.ijpara.2014.03.002

28. Siregar JE, Faust CL, Murdiyarso LS, Rosmanah L, Saepuloh U, Dobson AP, et al. Non ⍰ invasive surveillance for Plasmodium in reservoir macaque species. Malar J. 2015;1–8.

29. Kawai S, Megumi S, Kato-Hayashi N, Kishi H, Huffman MA, Maeno Y, et al. Detection of Plasmodium knowlesi DNA in the urine and faeces of a Japanese macaque (Macaca fuscata) over the course of an experimentally induced infection. Malar J. 2014;13(373):1–9.

30. Tan CH, Vythilingam I, Matusop A, Chan ST, Singh B. Bionomics of Anopheles latens in Kapit, Sarawak, Malaysian Borneo in relation to the transmission of zoonotic simian malaria parasite Plasmodium knowlesi. Malar J. 2008;7:1–8.

31. Vythilingam I, Noorazian YM, Huat TC, Jiram AI, Yusri YM, Azahari AH, et al. Plasmodium knowlesi in humans, macaques and mosquitoes in peninsular Malaysia. Parasit Vectors. 2008 Jan;1(1):1–10.

32. Jiram AI, Vythilingam I, NoorAzian YM, Yusof YM, Azahari AH, Fong M-Y. Entomologic investigation of Plasmodium knowlesi vectors in Kuala Lipis, Pahang, Malaysia. Malar J. 2012 Jan;11(213):1–10.

33. Hawkes F, Manin BO, Ng SH, Torr SJ, Drakeley C, Chua TH, et al. Evaluation of electric nets as means to sample mosquito vectors host-seeking on humans and primates. Parasit Vectors. 2017;10(1):338.

34. Hiwat H, Andriessen R, de Rijk M, Koenraadt CJM, Takken W. Carbon dioxide baited trap catches do not correlate with human landing collections of Anopheles aquasalis in Suriname. Mem Inst Oswaldo Cruz. 2011;106(3):360–4.

35. Hiwat H, de Rijk M, Andriessen R, Koenraadt CJM, Takken W. Evaluation of methods for sampling the malaria vector Anopheles darlingi (Diptera, Culicidae) in Suriname and the relation with its biting behavior. J Med Entomol. 2011;48(5):1039–46.

36. Sant’Ana DC, de Sá ILR, Sallum MAM. Effectiveness of Mosquito Magnet^®^ trap in rural areas in the southeastern tropical Atlantic Forest. Mem Inst Oswaldo Cruz. 2014; 109(8): 1021–9.

37. Xue R-D, Qualls W a, Kline DL, Zhao T-Y. Evaluation of lurex 3, octenol, and CO2 sachet as baits in Mosquito Magnet Pro traps against floodwater mosquitoes. J Am Mosq Control Assoc. 2010;26(3):344–5.

38. Chaves LSM, Laporta GZ, Sallum MAM. Effectiveness of mosquito magnet in preserved area on the coastal atlantic rainforest: Implication for entomological surveillance. J Med Entomol. 2014;51(5):915–24.

39. Vezenegho SB, Adde A, Gaborit P, Carinci R, Issaly J, Pommier de Santi V, et al. Mosquito magnet^®^ liberty plus trap baited with octenol confirmed best candidate for Anopheles surveillance and proved promising in predicting risk of malaria transmission in French Guiana. Malar J. 2014;13(1):384.

40. Li C-X, Dong Y-D, Zhang X-L, Chen C, Song S-P, Deng B, et al. Evaluation of Octenol and Lurex™ as baits in Mosquito Magnet^®^ Pro traps to collect vector mosquitoes in China. J Am Mosq Control Assoc. 2010;26(4):449–51.

41. Hing S. A survey of endoparasites in endangered Bornean elephants Elephas maximus borneensis in continuous and fragmented habitat. Imperial College London; 2012.

42. Boonratana R. The ecology and behaviour of the proboscis monkey (Nasalis larvatus) in the Lower Kinabatangan, Sabah. Mahidol University; 1994.

43. Estes JG, Othman N, Ismail S, Ancrenaz M, Goossens B, Ambu LN, et al. Quantity and Configuration of Available Elephant Habitat and Related Conservation Concerns in the Lower Kinabatangan Floodplain of Sabah, Malaysia. PLoS One. 2012;7(10).

44. Vythilingam I, Chan ST, Shanmugratnam C, Tanrang H, Chooi KH. The impact of development and malaria control activities on its vectors in the Kinabatangan area of Sabah, East Malaysia. Acta Trop. 2005;96(1):24–30.

45. Stark DJ, Vaughan IP, Evans LJ, Kler H, Goossens B. Combining drones and satellite tracking as an effective tool for informing policy change in riparian habitats: a proboscis monkey case study. Remote Sens Ecol Conserv. 2018;4(1):44–52.

46. Matsuda I, Otani Y, Bernard H, Wong A, Tuuga A. Primate survey in a bornean flooded forest: Evaluation of best approach and best timing. Mammal Study. 2016;41(2):101–6.

47. Goossens B, Setchell JM, Abulani A, Jalil MF, James SS, Aris SH, et al. A boat survey of primates in the Lower Kinabatangan Wildlife Sanctuary. Lower Kinabatangan scientific expedition. 2002. 37–45 p.

48. Rattanarithikul R, Harbach RE, Harrison BA, Panthusiri P, Jones JW, Coleman RE. Illustrated keys to the mosquitoes of Thailand II. Genera Culex and Lutzia. Southeast Asian J Trop Med Public Health. 2005;36:1–97.

49. Rattanarithikul R, Harrison B a., Panthusiri P, Coleman RE. Illustrated keys to the mosquitoes of Thailand. I. Background; geographic distribution; lists of genera, subgenera, and species; and a key to the genera. Southeast Asian J Trop Med Public Health. 2005;36:1–80.

50. Rattanarithikul R, Harrison BA, Harbach RE, Panthusiri P. Illustrated keys to the mosquitoes of Thailand IV. Anopheles. 2006;37:1–128.

51. Rattanarithikul R, Harrison B a., Panthusiri P, Peyton EL, Coleman RE. Illustrated keys to the mosquitoes of Thailand: III. Genera Aedeomyia, Ficalbia, Mimomyia, Hodgesia, Coquillettidia, Mansonia, and Uranotaenia. Southeast Asian J Trop Med Public Health. 2006;37:1–10.

52. Sallum MAM, Peyton EL, Harrison BA, Wilkerson RC. Revision of the Leucosphyrus group of Anopheles (Cellia) (Diptera, Culicidae). Rev Bras Entomol. 2005;49(SUPPL. 1):1–152.

53. Rahman WA, Hassan AA, Adanan CR, Rashid MR. The prevalence of Plasmodium falciparum and P. vivax in relation to Anopheles maculatus densities in a Malaysian village. Acta Trop. 1993;55(4):231–5.

54. Hawkes FM, Manin BO, Cooper A, Daim S, Homathevi R, Jelip J, et al. Vector compositions change across forested to deforested ecotones in emerging areas of zoonotic malaria transmission in Malaysia. Sci Rep. 2019;9(1):1–12.

55. Sriwichai P, Samung Y, Sumruayphol S, Kiattibutr K, Kumpitak C, Payakkapol A, et al. Natural human Plasmodium infections in major Anopheles mosquitoes in western Thailand. Parasites and Vectors [Internet]. 2016;9(1):1–9. Available from: http://dx.doi.org/10.1186/s13071-016-1295-x

56. Manguin S, Garros C, Dusfour I, Harbach RE, Coosemans M. Bionomics, taxonomy, and distribution of the major malaria vector taxa of Anopheles subgenus Cellia in Southeast Asia: An updated review. Infect Genet Evol. 2008;8(4):489–503.

57. Brown R, Chua TH, Fornace K, Drakeley C, Vythilingam I, Ferguson HM. Human exposure to zoonotic malaria vectors in village, farm and forest habitats in Sabah, Malaysian Borneo. PLoS Negl Trop Dis [Internet]. 2020;14(9):1–18. Available from: http://dx.doi.org/10.1371/journal.pntd.0008617

58. Lindén A, Mäntyniemi S. Using the negative binomial distribution to model overdispersion in ecological count data. Ecology. 2011;92(7):1414–21.

59. Rubio-Palis Y, Bevilacqua M, Medina DA, Moreno JE, Cárdenas L, Sánchez V, et al. Malaria entomological risk factors in relation to land cover in the Lower Caura River Basin, Venezuela. Mem Inst Oswaldo Cruz. 2013;108(2):220–8.

60. Dekel A, Pitts RJ, Yakir E, Bohbot JD. Evolutionarily conserved odorant receptor function questions ecological context of octenol role in mosquitoes. Sci Rep [Internet]. 2016;6(November):1–7. Available from: http://dx.doi.org/10.1038/srep37330

61. Fenske JD, Paulson SE. Human breath emissions of VOCs. J Air Waste Manag Assoc. 1999;49(5):594–8.

62. McCann RS, Messina JP, MacFarlane DW, Bayoh MN, Gimnig JE, Giorgi E, et al. Explaining variation in adult Anopheles indoor resting abundance: The relative effects of larval habitat proximity and insecticide-treated bed net use. Malar J. 2017;16(1):1–14.

63. Kaindoa EW, Mkandawile G, Ligamba G, Kelly-Hope LA, Okumu FO. Correlations between household occupancy and malaria vector biting risk in rural Tanzanian villages: Implications for high-resolution spatial targeting of control interventions. Malar J. 2016;15(1):1–12.

64. Goossens B, Ambu LN. Sabah Wildlife Department and 10 years of research: Towards a better conservation of Sabah’s wildlife. J Oil Palm Environ. 2012;3(5):38–51.

65. Collins WE, Sullivan JS, Galland GG, Nace D, Williams A, Williams T, et al. Isolates of Plasmodium inui adapted to Macaca mulatta monkeys and laboratory-reared Anopheline mosquitoes for experimental study. J Parasitol. 2007;93(5):1061–9.

66. Vythilingam I, Lee K-S. Plasmodium knowlesi: Emergent Human Malaria in Southeast Asia. In: Lim YAL, Vythilingam I, editors. Parasites and their vectors. Springer; 2013. p. 131–54.

67. Akter R, Vythilingam I, Khaw LT, Qvist R, Lim YAL, Sitam FT, et al. Simian malaria in wild macaques: first report from Hulu Selangor district, Selangor, Malaysia. Malar J. 2015;14(1):1–9.

68. Bt K, Kadir A, Cliff P, Divis S, Shuaisah D, Awang B, et al. Zoonotic Malaria Parasites Among Non-Human Primates in Sarawak, Malaysian Borneo. 2013;2.

69. Lee KS, Divis PCS, Zakaria SK, Matusop A, Julin RA, Conway DJ, et al. Plasmodium knowlesi: Reservoir hosts and tracking the emergence in humans and macaques. PLoS Pathog. 2011;7(4).

70. Loy DE, Rubel MA, Avitto AN, Liu W, Li Y, Learn GH, et al. Investigating zoonotic infection barriers to ape Plasmodium parasites using faecal DNA analysis. Int J Parasitol [Internet]. 2018;48(7):531–42. Available from: https://doi.org/10.1016/j.ijpara.2017.12.002

71. Zhang X, Kadir KA, Quintanilla-Zariñan LF, Villano J, Houghton P, Du H, et al. Distribution and prevalence of malaria parasites among long-tailed macaques (Macaca fascicularis) in regional populations across Southeast Asia. Malar J. 2016;15(1):1–8.

72. Chua TH, Manin BO, Daim S, Vythilingam I, Drakeley C. Phylogenetic analysis of simian Plasmodium spp. infecting Anopheles balabacensis Baisas in Sabah, Malaysia. PLoS Negl Trop Dis. 2017;11(10):1–13.

73. Wong ML, Chua TH, Leong CS, Khaw LT, Fornace K, Wan-Sulaiman WY, et al. Seasonal and Spatial Dynamics of the Primary Vector of Plasmodium knowlesi within a Major Transmission Focus in Sabah, Malaysia. PLoS Negl Trop Dis. 2015;9(10):1–15.

