## Supplementary figures and images for "Measuring the exposure of primate reservoir hosts to mosquito vectors in Malaysian Borneo"

### Figure S1

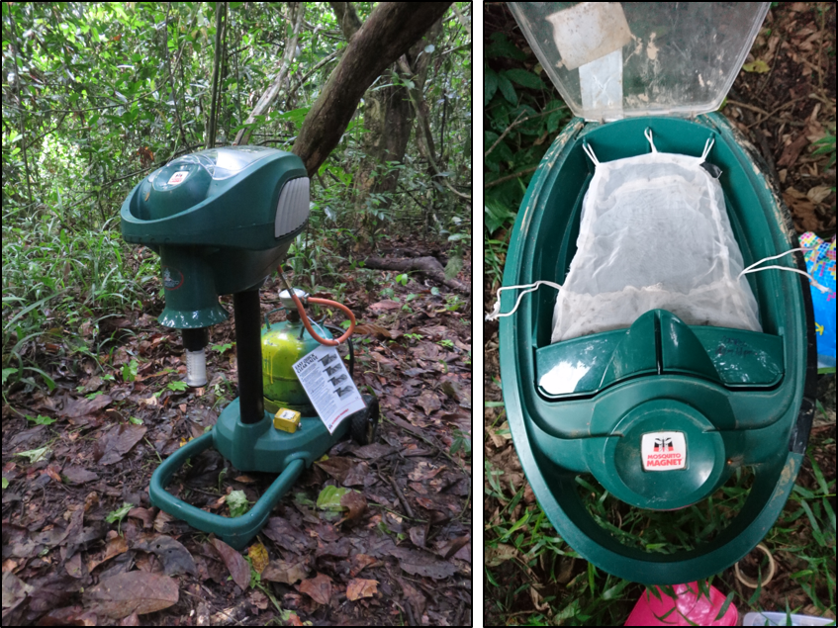

### Figure S2

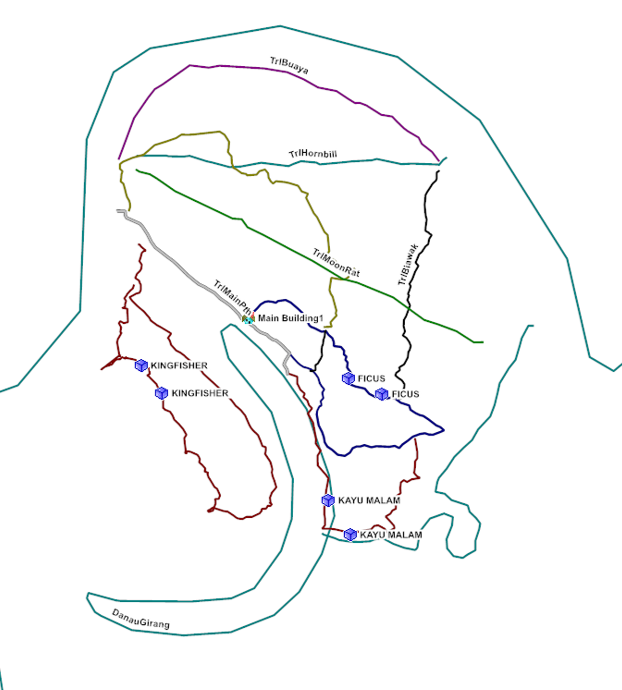

### Figure S3

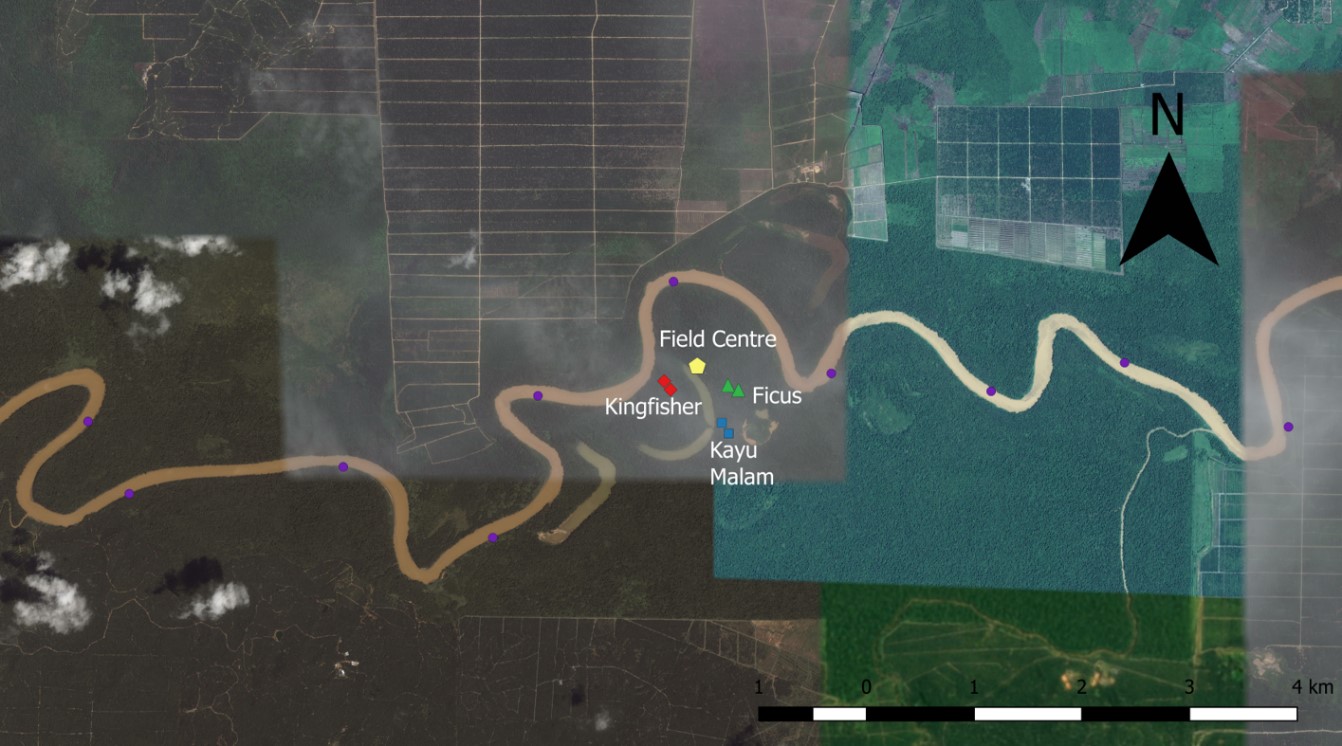

### Figure S4

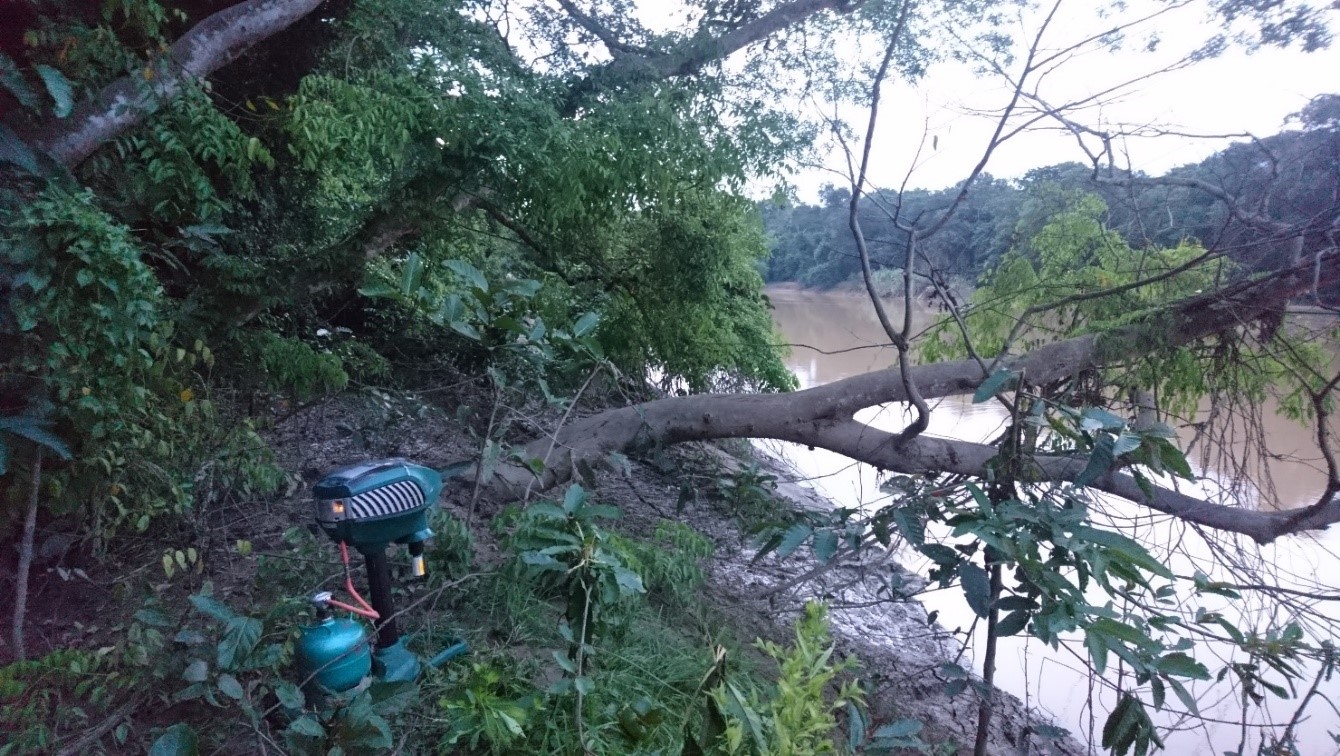

### Figure S5

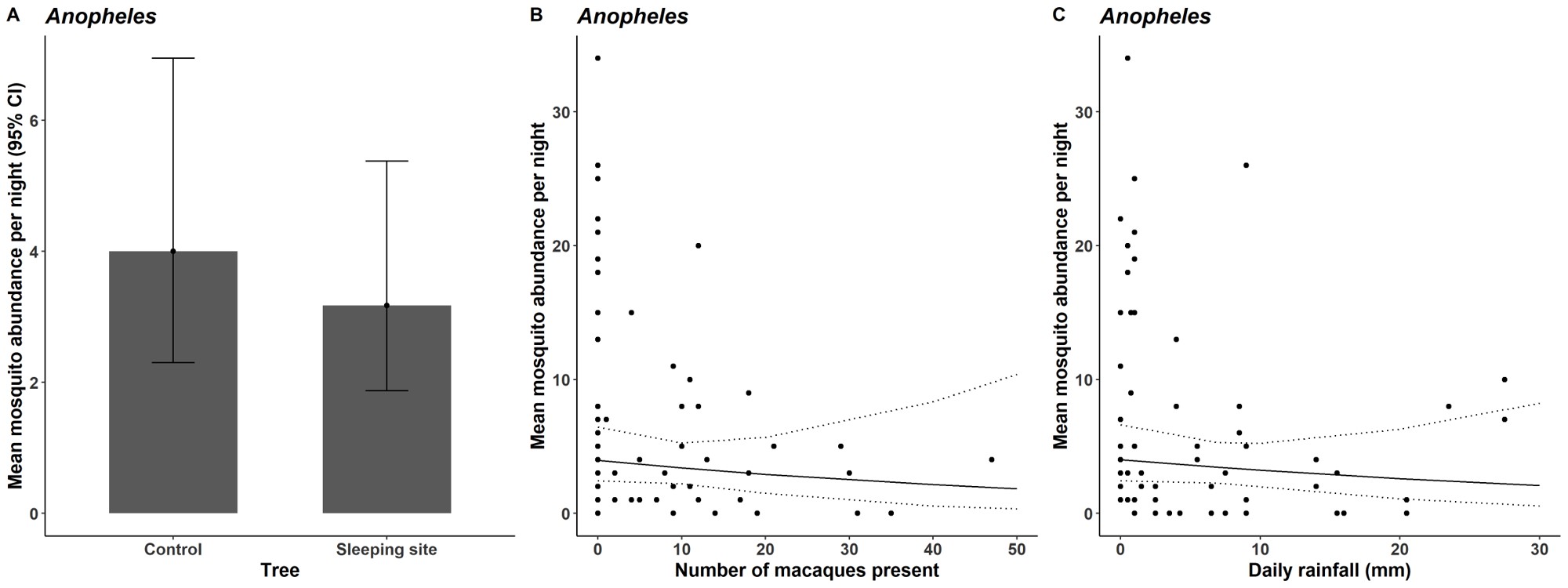

### Figure S6

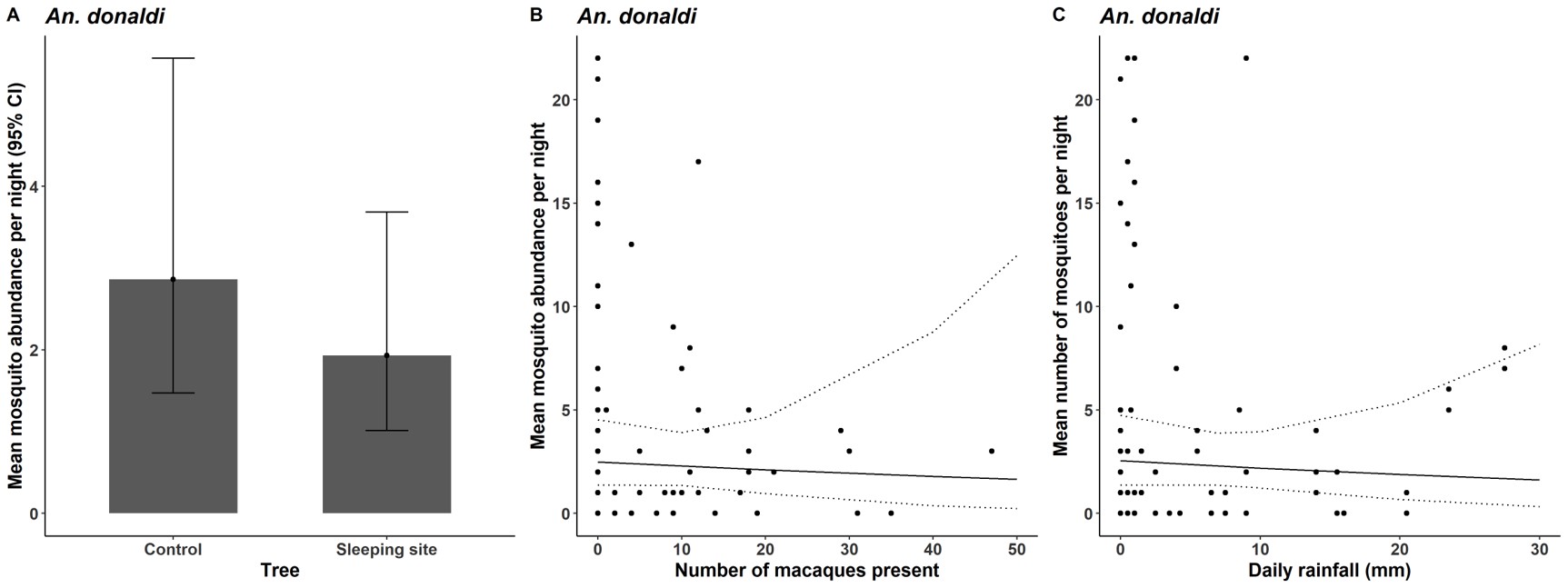

### Figure S7

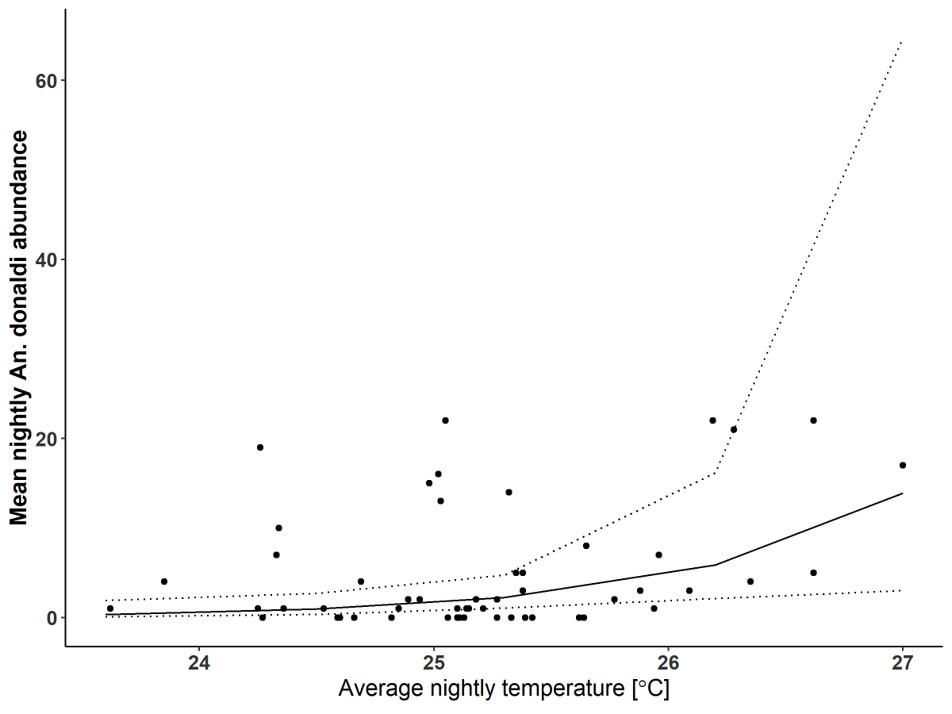
