## Supplementary methods for "Measuring the exposure of primate reservoir hosts to mosquito vectors in Malaysian Borneo"

**Methods S1. Macaque faecal sample collection.**

Two people conducted a twenty-minute search of the forest floor and canopy (within reach). Efforts were made to ensure each faecal sample corresponded to a different individual based on variation in the colour, consistency and distance of sample from other faeces. As many samples as possible were taken within these limitations, however there were some cases where no faeces could be found. Wearing gloves and a N95 disposable respirator face-mask, a 2 cm^3^ faecal sample was deposited into a 50 ml falcon tube and sealed in a zip-lock plastic bag. Tubes were immediately transported to the laboratory and filled with RNAlater solution. The sample was homogenized immediately using a sterile chopstick until completely broken. Tubes were stored at -20˚C until further processing.

**Methods S2. PCR screening of DNA extracted from macaque stool samples.**

Nested PCRs were conducted to screen samples for *Plasmodium* DNA firstly using the method of Siregar (1), which identifies DNA of any species within the *Plasmodium* genus. 2μl of genomic DNA was subjected to an outer amplification reaction with 0.4 μM of each of the SSU-rRNA *Plasmodium* genus specific primers PfF4595 (GATTACAGCTCCCAAGCAAAC) and PfR5019 (GTTTAGCCAGGAAGTCAGCGTC), 100 μM dNTPs, and 0.5 U Phusion High Fidelity DNA polymerase (New England Biolabs M0530) with 1 x High Fidelity buffer in a total volume of 25μl. The nested reaction was identical except that only 1µl PCR product from nest 1 was used for nest 2. PCR conditions were: initial denaturation at 94 ^o^C for 5 min; followed by 30 cycles of 94 ^o^C for 15 seconds, 60 ^o^C annealing for 15 seconds and 72 ^o^C for 45 seconds; and a final extension at 72 ^o^C for 5 minutes. 1µl of PCR product from nest 1 was used for nest 2. Nest 2 was an exact replicate of nest 1 but with 16.5 µl dH_2_O. PCR products were run on 3 % agarose gel in 0.5x TAE buffer. Samples with a 424 bp band were positive for *Plasmodium* genus.

After initial screening for *Plasmodium*, all positive samples went through another round of PCR to test for the specific presence of *P. knowlesi* following the method of Kawai *et al* (2)*.* Here 2μl of genomic DNA was subjected to an outer amplification reaction with 0.5 μM of each of the cytochrome b *P. knowlesi* primers PCBF (ATGCTTTATTATGGATTGGATGTC) and PCBRed (ACATAATTATAACCTTACGGTCTG), 100 μM dNTPs, and 0.5 U Phusion High Fidelity DNA polymerase (New England Biolabs M0530) with 1 x High Fidelity buffer in a total volume of 25μl. The nested reaction was identical except for substitution of the primers with PkCBF (TATTCTTCTTTAGTGGATTATTTA) and PkCBRed (GTATTGTTCTAATCAGTGTA), and the use of 1 μl of outer PCR product instead of genomic DNA. PCR conditions were initial denaturation at 98 ^o^C for 1 min; followed by 35 cycles of 98 ^o^C for 10 seconds, 50 ^o^C annealing for 30 seconds and 72 ^o^C for 30 seconds; and a final extension at 72 ^o^C for 5 minutes. PCR products were run on 3 % agarose gel in 0.5x TAE buffer. A 131bp band indicated that the sample was positive for *P. knowlesi* DNA.

1.	Siregar JE, Faust CL, Murdiyarso LS, Rosmanah L, Saepuloh U, Dobson AP, et al. Non ‑ invasive surveillance for Plasmodium in reservoir macaque species. Malar J. 2015;1–8.

2.	Kawai S, Megumi S, Kato-Hayashi N, Kishi H, Huffman MA, Maeno Y, et al. Detection of Plasmodium knowlesi DNA in the urine and faeces of a Japanese macaque (Macaca fuscata) over the course of an experimentally induced infection. Malar J. 2014;13(373):1–9.
