## Supplementary tables for "Measuring the exposure of primate reservoir hosts to mosquito vectors in Malaysian Borneo"

| Mosquito genus/ subgenus/ species | Human-landing catch (HLC) | Mosquito Magnet Independence Trap (MMIT) |
| --- | --- | --- |
| ***Aedes*** | **87** | **151** |
| *Aedimorphus* | 2 | 8 |
| *Am. Caecus* | 2 | 2 |
| *Ayurakitia* | 0 | 1 |
| *Downsiomyia* | 1 | 0 |
| *Edwardsaedes* | 3 | 1 |
| *Finlaya* | 2 | 12 |
| *Paraedes* | 67 | 81 |
| *Pr. Ostentato* | 10 | 0 |
| *Ochlerotatus* | 3 | 10 |
| Unknown *Aedes* spp. | 9 | 38 |
| ***Anopheles*** | **36** | **62** |
| *An. balabacensis* | 2 | 5 |
| *An. barbirostris* | 0 | 2 |
| *An. barbirostris/donaldi* | 1 | 5 |
| *An. barbumbrosus* | 0 | 2 |
| *An. cellia* subgenus | 0 | 1 |
| *An. donaldi* | 27 | 40 |
| *An. kochi* | 0 | 1 |
| *An. montanus* | 2 | 1 |
| *An. roperi* | 1 | 1 |
| *An. tesselatus* | 3 | 2 |
| Unknown *Anopheles* spp. | 0 | 2 |
| ***Coquillettidia*** | **2** | **23** |
| ***Culex*** | **278** | **532** |
| *Cx. brevipalpis/phangngae* | 1 | 0 |
| *Cx. foliates* | 21 | 5 |
| *Cx. fuscocephala* | 13 | 13 |
| *Cx. gelidus* | 1 | 6 |
| *Cx. hutchisoni* | 25 | 37 |
| *Cx. malayi* | 3 | 0 |
| *Cx. perplexus/Cx. whitei* | 0 | 2 |
| *Cx. pseudosiniensis* | 9 | 5 |
| *Cx. quinquefasciatus* | 8 | 4 |
| *Cx. siniensis* | 1 | 2 |
| *Cx. sitiens* | 5 | 1 |
| *Cx. tenuipalpis sub* | 47 | 97 |
| *Cx. vishnui/ pseudovishnui* | 31 | 179 |
| *Cx. whitmorei* | 78 | 30 |
| Unknown *Culex* spp. | 35 | 151 |

Table S1. Mosquitoes caught by Mosquito Magnet Independence Trap (MMIT) and human-landing catch (HLC) over ten nights of trap comparison study in Lower Kinabatangan Wildlife Sanctuary, Sabah.

Table S1 continued on next page

Table S1 continued. Mosquitoes caught by Mosquito Magnet Independence Trap (MMIT) and human-landing catch (HLC) over ten nights of trap comparison study in Lower Kinabatangan Wildlife Sanctuary, Sabah.

| Mosquito genus/ subgenus/ species | Human-landing catch (HLC) | Mosquito Magnet Independence Trap (MMIT) |
| --- | --- | --- |
| ***Mansonia*** | **396** | **300** |
| *Ma. annulate* | 122 | 98 |
| *Ma. annulifera* | 3 | 3 |
| *Ma. bonnae* | 13 | 3 |
| *Ma. dives* | 10 | 15 |
| *Ma. dives/ bonnae* | 105 | 63 |
| *Ma. Indiana* | 86 | 63 |
| *Ma. uniformis* | 48 | 29 |
| Unknown Mansonia spp. | 9 | 26 |
| ***Orthopodomyia*** | **3** | **3** |
| ***Uranotaenia*** | **4** | **9** |
| *Ur. Longirostris* | 0 | 2 |
| *Ur.* species 3 | 1 | 2 |
| Unknown *Uranotaenia* spp. | 3 | 5 |
| ***Verrallina*** | **237** | **746** |
| Unknown genera | 3 | 11 |
| **Total** | **1049** | **1846** |

Table S2. Mosquitoes caught with Mosquito Magnet Independence Trap (MMIT) at trees with and without sleeping macaques (control trees) within the Lower Kinabatangan Wildlife Sanctuary, Sabah.

| Mosquito genus/ subgenus/ species | Macaque sleeping sites (34 nights) | Control trees  (33 nights) |
| --- | --- | --- |
| ***Aedes*** | **193** | **169** |
| *Ae. laniger* | 1 | 0 |
| *Aedimorphus* | 0 | 1 |
| *Ayurakitia* | 0 | 1 |
| *Edwardsaedes* | 29 | 1 |
| *Finlaya* | 1 | 0 |
| *Paraedes* | 142 | 130 |
| *Ochlerotatus* | 15 | 22 |
| *Scutomyia albolineata* | 1 | 0 |
| *Stegomyia* | 0 | 1 |
| Unknown *Aedes* spp. | 4 | 13 |
| ***Anopheles*** | **250** | **476** |
| *An. balabacensis* | 13 | 2 |
| *Barbirostris gp* | 122 | 251 |
| *An. barbirostris* | 0 | 2 |
| *An. donaldi* | 106 | 211 |
| *An. epiroticus* | 1 | 0 |
| *An. gigas* | 1 | 0 |
| *An. kochi* | 0 | 0 |
| *An. montanus* | 2 | 2 |
| *An. roperi* | 0 | 1 |
| *An. tesselatus* | 2 | 0 |
| *An. umbrosus gp* | 1 | 2 |
| Unknown *Anopheles* spp. | 2 | 5 |
| ***Coquillettidia*** | **37** | **23** |
| *Coq. Nigrosignata* | 19 | 18 |
| Unknown *Coquillettidia* spp. | 18 | 5 |
| ***Culex*** | **1581** | **1213** |
| *Cx. baileyi* | 0 | 1 |
| *Cx. brevipalpis/phangngae* | 23 | 2 |
| *Cx. cinctellus* | 21 | 13 |
| *Cx. foliatus* | 4 | 3 |
| *Cx. fuscocephala* | 0 | 1 |
| *Cx. gelidus* | 42 | 27 |
| *Cx. hutchisoni* | 399 | 350 |
| *Cx. infantulus* | 0 | 1 |
| *Cx. infula* | 1 | 8 |
| *Cx. mammifer/ wilfredi* | 2 | 26 |
| *Cx. nigropunctatus* | 189 | 132 |
| *Cx. pseudosinensis* | 43 | 21 |
| *Cx. pseudovishnui/vishnui* | 277 | 209 |
| *Cx. quinquefasciatus* | 2 | 1 |
| *Cx. scanloni* | 1 | 0 |
| *Cx. siniensis* | 5 | 1 |

Table S2 continued on next page

Table S2 continued. Mosquitoes caught with Mosquito Magnet Independence Trap (MMIT) at trees with and without sleeping macaques (control trees) within the Lower Kinabatangan Wildlife Sanctuary, Sabah.

| Mosquito genus/ subgenus/ species | Macaque sleeping sites | Control trees |
| --- | --- | --- |
| *Cx. sitiens* | 24 | 16 |
| *Cx. tenuipalpis* | 277 | 198 |
| *Cx. whitmorei* | 133 | 53 |
| *Cx. whitmorei/gelidus* | 0 | 3 |
| *Cx. whitei* | 0 | 1 |
| Unknown *Culex* spp. | 138 | 146 |
| ***Mansonia*** | **3151** | **3036** |
| *Ma. annulate* | 468 | 467 |
| *Ma. annulifera* | 254 | 265 |
| *Ma. bonnae* | 220 | 153 |
| *Ma. dives* | 695 | 416 |
| *Ma. dives/bonnae* | 186 | 232 |
| *Ma. Indiana* | 300 | 588 |
| *Ma. uniformis* | 191 | 148 |
| Unknown Mansonia spp. | 837 | 767 |
| ***Orthopodomyia*** | **2** | **4** |
| ***Uranotaenia*** | **48** | **63** |
| *Uranotaenia* | 11 | 19 |
| *Pseudoficalbia* | 30 | 33 |
| Unknown *Uranotaenia* spp. | 7 | 11 |
| ***Verrallina*** | **631** | **523** |
| **Total** | **5893** | **5507** |
